# *Massilia varians* P2-4, A Potential Biocontrol Agent against Pathogenic *Pseudomonas aeruginosa* in *Eriocheir sinensis*

**DOI:** 10.64898/2026.05.13.725027

**Authors:** Yiyao Liu, Yueqi Yang, Meiling Liu, Sijia Chen, Haipeng Cao, Chunlei Gai, Weidong Ye

## Abstract

*Pseudomonas aeruginosa* is a clinically significant bacterial pathogen that poses a serious threat to aquaculture. However, there are limited information on *Massilia* isolates against pathogenic *P. aeruginosa* in aquaculture. In the present study, a facultative predator, *M. varians* isolate P2-4, was isolated from aquaculture sediment using Chinese mitten crab *Eriocheir sinensis*-pathogenic *P. aeruginosa* as the prey bacterium, and its genomic feature, bacteriolysis-related genes, safety, bacteriolytic spectrum, and *in vitro* and *in vivo* antibacterial effects against pathogenic *P. aeruginosa* in *E. sinensis* were further characterized. Isolate P2-4 consisted of one chromosome and one plasmid (with a total of 75 tRNAs, 7 5S rRNAs, 7 16S rRNAs, 7 23S rRNAs, 34 sRNAs, 5,238 coding genes, 20 genomic islands, 1 prophage, 23 insertion sequences, and 102 repeat sequences), and harbored 19 bacteriolysis-related genes (*pilA, pilB, pilC, pilD, pilF, pilG, pilH, pilM, pilO, pilP, pilQ, pilS, pilR, pilT, mltA, mltB, mltC, mltD*, and *dacB*) associated with cellular motility and cell wall lysis. In addition, the isolate carried no virulence genes, was unable to produce haemolysin, hydrogen sulfide, nitrite and ammonia, and avirulent in *E. sinensis* with a 7-day acute intraperitoneal LD_50_ value of above 5.0 × 10^8^ CFU/mL. Furthermore, the isolate possessed a wide bacteriolytic spectrum against pathogenic *Shewanella algae, Aeromonas caviae, A. hydrophila*, and *Photobacterium damselae* besides *P. aeruginosa*, exhibited bacteriolysis rates of 99.35% to 99.99% towards the pathogenic *P. aeruginosa* at 1.0×10^3^ to 1.0×10□ CFU/mL, and displayed relative percentage survivals of 42.31% to 73.08% against *P. aeruginosa* infection in *E. sinensis* at doses of 6.0 × 10^3^ to 6.0 × 10^5^ CFU/g diet. To our knowledge, this study for the first time demonstrates a *M. varians* strain as a potential biocontrol agent against pathogenic *P. aeruginosa* in aquaculture.

## 1 Introduction

*Pseudomonas aeruginosa* is a Gram-negative zoonotic bacterium that is widely distributed in diverse ecological environments, including soil, water, and sewage (Liao et al., 2022). This bacterium can not only cause chronic obstructive pulmonary disease, malignant external otitis, endophthalmitis, and meningitis (Fong and Tomkins, 1985; Rubin and Yu, 1988; Eifrig et al., 2003; Murphy et al., 2008), but result in gill necrosis in *Oziotelphusa senex senex*, haemorrhagic septicaemia in *Oreochromis niloticus*, and shell erosion disease in *Bellamya aeruginosa* (Devakumar et al., 2013; El-Tarabili et al., 2023; Zhang et al., 2025). Currently, the primary method for controlling *P. aeruginosa* infections in aquaculture relies heavily on antibiotics (Ali et al., 2021; Kholil et al., 2015). However, frequent use of antibiotics increases the risk of antimicrobial resistance and may disrupt the aquatic ecological balance (Frascaroli et al., 2024; Pepi and Focardi, 2021). Thus, environmental-friendly control agents should be urgently exploited to mitigate *P. aeruginosa* infections in aquaculture.

Predatory bacteria are a diverse group of Gram-negative microorganisms that have the capacity to prey on or inhibit other bacteria by secreting lytic proteins and lysing cell walls (Novick, 2021; Kamada et al., 2023; Mohsenipour et al., 2024). They are grouped into facultative predator (prey-independent growth on organic nutrients) and obligate predator (predatory growth on living prey cells) (Bauer and Forchhammer, 2021). Currently, obligate predator such as *Bdellovibrio* and *Halobacteriovorax* exhibit remarkable inhibition against *Acinetobacter venetianus, Aeromonas hydrophila, A. veronii, Proteus penneri, Vibrio cholerae, V. parahaemolyticus*, and *V. vulnificus* pathogens in aquaculture (Cao et al., 2012; Cao et al., 2014a; Cao et al., 2014b; Cao et al., 2015; Cao et al., 2019; Huang et al., 2020; Ottaviani et al., 2018), and provide significant protection of aquaculture animals against *A. veronii* and *Shewanella putrefaciens* infections (Liu et al., 2022; Liu et al., 2024). However, scarce information is available on facultative predator against *P. aeruginosa* in aquaculture.

*Massilia* is a genus of Gram-negative bacteria with a broad distribution across diverse environments, including soil, aquatic systems, and the surfaces of various organisms (Xu et al., 2023). Although the role of *Massilia* remains controversial due to a few isolates are occasionally associated with human septicemia and osteomyelitis (Lindquist et al., 2003), it is still considered as a potential probiotic and exhibits excellent antibacterial effects (Yang et al., 2019). For example, *M. rhizosphaerae* NEAU-GH312^T^, *M. pseudoviolaceinigra* P3689^T^, and *M. scottii* P5043^T^ show strong antibacterial activity against *Ralstonia solanacearum, Curtobacterium flacumfaciens*, and *Paenarthrobacter ilicis* (Li et al., 2021; Sedláček et al., 2025). Nevertheless, there is currently no documented evidence suggesting *Massilia* strains as facultative predators against *P. aeruginosa* in aquaculture.

Chinese mitten crab *Eriocheir sinensis* is an economically important aquatic product in East and Southeast Asian countries (Agbekpornu et al., 2018), However, the aquaculture of this crab has recently been severely by *P. aeruginosa* (He et al., 2026). In this study, a facultative predator, *M. varians* isolate P2-4, against *E. sinensis*-pathogenic *P. aeruginosa* was isolated from aquaculture sediment, and its genomic feature, bacteriolysis-related genes, safety, bacteriolytic spectrum, and *in vitro* and *in vivo* antibacterial effects against the pathogenic *P. aeruginosa* were further characterized. To our knowledge, this is the first study to identify a *M. varians* strain as a potential biocontrol agent against pathogenic *P. aeruginosa* in aquaculture.

## 2 Materials and methods

### 2.1 Animal ethics

All animal experiments and experimental protocols were approved by the Institutional Animal Care and Use Ethics Committee of Shanghai Ocean University under approval no. SHOU-DW-2025-050.

### 2.2 Isolation of facultative predatory isolate

The isolation of facultative predatory isolate was carried out according to Wang (2020). Prior to the isolation, pathogenic *P. aeruginosa* strain HX-1, isolated from diseased *E. sinensis* suffering from hepatopancreas necrosis disease and identified through genome sequencing and phenotypic methods (He et al., 2026), was provided by the National Pathogen Collection Center for Aquatic Animals, Shanghai, China. The suspension of *P. aeruginosa* strain HX-1 was freshly obtained from 12h of strain HX-1 culture in nutrient broth (NB, Sinopharm Chemical Reagent Co., Ltd., Shanghai, China) by centrifuging (4000 r/min, 10 min) and suspending with normal saline (Wingender et al., 2001), and was finally quantified as 5.0×10^9^ CFU/mL by counting colony forming units (CFU) on nutrient agar (NA, Qingdao Hope Bio-Technology Co., Ltd., Shandong, China) plates from a series of ten-fold dilutions in sterile normal saline (Rodgers et al., 2023). The aquaculture sediment (1.0 g), collected from an aquaculture farm in Shandong, China, was suspended in 9.0 mL of sterile normal saline and vortexed at 30°C for 30 min to achieve a mixture. The mixture was then subjected to a series of ten-fold serial dilutions ranging from 10^−2^, 10^−3^, 10^−4^, 10^−5^, and 10^−6^ in sterile normal saline. Subsequently, these dilutions were examined for the presence of plaques using *P. aeruginosa* strain HX-1 as the prey bacterium by double-layer agar plate method (Ottaviani et al., 2020). Briefly, 100 μL of the dilution and 200 μL of the *P. aeruginosa* suspension were added to 10 mL of 0.6% agar melted and kept at 55 °C, mixed and poured onto a pre-prepared 1.5% agar plate until solid. The plaques were observed carefully after the incubation of the double-layer plates at 30 °C for 5 days. Aliquots (0.2 mL) of the dilutions with plaque present were then spread onto NA plates and incubated at 30°C for 24h. Individual bacterial colonies were selected using a flame-sterilized inoculating loop and further purified through restreaking and subculturing on NA plates under the same incubation conditions. The facultatively predatory isolate was finally verified by plaque-forming ability examination of pure isolates via double-layer agar plate method (Ottaviani et al., 2020), using *P. aeruginosa* strain HX-1 as the prey bacterium. Facultatively predatory isolate was inoculated onto the NA slant and kept at 4°C following the incubation at 30°C for 24h.

### 2.3 Identification of facultatively predatory isolate

#### 2.3.1 16S rRNA gene sequencing analysis

The genomic DNA of the isolate was extracted using a TIANamp bacteria DNA kit (Tiangen Biotech Co., Ltd., Beijing, China). The 16S rRNA gene of the isolate was amplified via PCR with universal primers 27F (5’-AGAGTTTGATCCTGGCTCAG-3’) and 1492R (5’-GGTTACCTTGTTACGACTT-3’) according to the method outlined by Tian et al. (2023). The PCR reaction mixture consisted of 10 µM of each primer (1 µL), 2×Taq Master mix (Vazyme, Nanjing, China) (20 µL), 20 ng/µL of template DNA (1 µL), and ddH_2_O (17 µL). The PCR amplification conditions included initial denaturation at 95°C for 3 min, followed by 35 cycles of denaturation at 95°C for 1 min, annealing at 60°C for 1 min, and extension at 72°C for 1 min, with a final extension at 72°C for 10 min. The resultant PCR product was subjected to 1.0% (w/v) agarose gel electrophoresis, and sequenced by Shanghai MAP Biotech. Co., Ltd., Shanghai, China. The obtained sequence was subjected to homology analysis using the Basic Local Alignment Search Tool (BLAST) (https://blast.ncbi.nlm.nih.gov/Blast.cgi) against the GenBank database. Phylogenetic analysis was conducted using neighbor-joining method in MEGA 11.0 software (Pennsylvania State University, Pennsylvania, USA).

#### 2.3.2 Genomic sequencing analysis

Whole genome sequencing of the isolate was conducted using the Illumina HiSeq and PacBio platforms (Shanghai Majorbio Biopharm Technology Co., Ltd., Shanghai, China) according to Li et al. (2023). The generation of sequencing reads was followed by their assembly using SOAPdenovo2 and CANU software (Luo et al., 2012; Koren et al., 2017). Concurrently, the construction of circular genome maps was undertaken with the use of Circos (v.0.69) (Krzywinski et al., 2009). A comprehensive analysis of various genomic features, including Genome size (bp), GC average content, gene average length, repeated regions, number of chromosomes and plasmids, tRNA, rRNA, sRNA, coding sequences (CDS), tandem repeats, prophages, genomic islands (GI), and insertion sequences (IS), were predicted using tRNAscan-SE (v.2.0), Barrnap (v.0.8), Infernal, Glimmer (v.3.02), GeneMarkS, Tandem Repeats Finder (v.4.07), Phage Finder, IslandPath-DIMOB (v.1.0.0), and ISEScan software (Benson, 1999; Besemer and Borodovsky, 2005; Chan and Lowe, 2019; Fouts, 2006; Xie and Tang, 2017). Overall genome-relatedness indices between the isolate and closely related bacterial species, including the average nucleotide identity (ANI) and digital DNA–DNA hybridization (dDDH), were calculated using MUMmer (v.3.23) and Genome-to-Genome Distance Calculator (v.3.0) software (Ribeiro et al., 2021), at the 95% ANI and 70% dDDH thresholds for species delineation (Candeliere et al., 2021).

#### 2.3.3 Phenotypic identification

Following careful examinations of Gram stain and growth on thiosulfate-citrate-bile salts-sucrose agar (TCBS, Qingdao Hope Bio-Technology Co., Ltd., Shandong, China), the phenotypic characterization of the isolate was conducted in triplicate using API 20NE test strips (bioMérieux, Marcy-l’Étoile, France) in accordance with the manufacturer’s instructions (Roxo et al., 2025). The phenotypic traits of the isolate were compared with those as previously described (Cho et al., 2017; Chaudhary and Kim., 2017; Orthova et al.,2015; Weon et al., 2010).

### 2.4 Virulence and bacteriolysis-related gene assay

The BLAST analysis of the isolate genome (obtained in section 2.3.2) was carried out against the virulence factors database (VFDB, http://www.mgc.ac.cn/VFs/) to identify virulence genes (Liu et al., 2019). Clusters of Orthologous Genes (COG) and Kyoto Encyclopedia of Genes and Genomes (KEGG) analyses in the isolate genome was also conducted to identify the bacteriolysis-related genes described by Kanehisa and Goto. (2000), using BLAST2GO (v.1.0) and DIAMOND blastp (v.2.0.14) software (Tatusov et al., 2000; Kanehisa, 2000). The cutoff values were set at E-value ≤ 1e^−5^, identity ≥ 90% and coverage ≥ 90% (Wang et al., 2021).

### 2.5 Toxic metabolite production assay

#### 2.5.1 Haemolysin production assay

The haemolysin production of the isolate was examined in triplicate with rabbit blood agar (RBA) plates according to Gai et al. (2023). Briefly, the isolate was inoculated onto the RBA plates (Qingdao Hope Bio-Technology Co., Ltd., Shandong, China) and incubated at 30°C for 48h to observe the presence of zones surrounding bacterial colony. The presence of an incomplete transparent zone or a clear colorless zone surrounding bacterial colonies has been recorded as indicative of the presence of α-haemolysin or β-haemolysin (Castro-Escarpulli et al., 2003). *Rhodobacter azotoformans* strain SY5, a safe strain for *E. sinensis* isolated from aquaculture sediment and identified through molecular and phenotypic methods (Cao et al., 2022; Cao et al., 2023), was used as the control.

#### 2.5.2 Hydrogen sulfide, nitrite, and ammonia production assay

The isolate was examined in triplicate for the production of hydrogen sulfide, nitrite and ammonia using microbiochemical identification tubes as recommended by Kang et al. (2019). Briefly, the isolate was inoculated into the hydrogen sulfide, nitrite, and ammonia biochemical identification tubes following the manufacturer’s guidelines (Qingdao Hope Bio-Technology Co., Ltd., Shandong, China), and incubated at 37°C for 24 h. The hydrogen sulfide, nitrite, and ammonia production of the isolate is determined in terms of the color of tubes following the manufacturer’s instructions. *R. azotoformans* strain SY5 was used as the control.

### 2.6 Bacteriolytic spectrum assay

Prior to this assay, the isolate was inoculated into NB for 24h of culture at 30°C, harvested by centrifugation at 4000 r/min for 10 min, and the resulting pellet was suspended in normal saline to prepare the isolate suspension. Thirteen pathogenic strains (*P. aeruginosa* strains FJ1-2, FJ1-5, FJ1-9, FJ3-2, and PX3-3, *S. algae* strains RZ2-2, RZ3-4, and RZ2-1, *A. caviae* strains PX2-5 and PX3-1, *A. hydrophila* strains PX2-8 and PX2-6, and *Photobacterium damselae* strain YJ-1) were obtained from National Pathogen Collection Center for Aquatic Animals, Shanghai, China. The suspensions of these pathogenic strains were freshly prepared as described in section 2.2. The suspensions of the isolate and pathogenic strains were quantified through counting CFU on NA plates from a series of ten-fold dilutions in sterile normal saline. The bacteriolysis of the isolate was examined in triplicate against the thirteen pathogenic strains using double-layer agar plate method (Ottaviani et al., 2020). Briefly, 50 μL of the isolate suspension (1.0×10^6^ CFU/mL) and 100 μL of the pathogenic strain suspension (5.0×10^9^ CFU/mL) were added to 10 mL of 0.6% agar melted and kept at 55 °C, mixed and poured onto a pre-prepared 1.5% agar plate until solid. Following incubation at 30°C for 48h, the presence of plaques on the double-layer agar plates was observed, and the number of plaque forming units (PFU) was recorded. The bacteriolysis of the isolate against *P. aeruginosa* strain HX-1 (5.0×10^9^ CFU/mL) was used as the control.

### 2.7 In vitro antibacterial effect against *P. aeruginosa* assay

The *in vitro* antibacterial effect of the isolate against *P. aeruginosa* was examined in twelve 250□mL-glass flasks, which were divided into one control and three treatment groups (three flasks per group). Prior to this assay, the suspensions of the isolate and *P. aeruginosa* strain HX-1 were freshly prepared as described in sections 2.2 and 2.6. Following quantification through counting CFU on NA plates from a series of ten-fold dilutions in sterile normal saline, the suspensions of the isolate and *P. aeruginosa* were respectively added to the treatment groups of flasks, and the filtered farming water, prepared and autoclaved according to Cao et al. (2019), was then immediately supplemented to the treatment groups of flasks to a final volume of 200 mL to achieve 1.0 × 10^3^, 1.0 × 10□, and 1.0 × 10□ CFU/mL of the isolate and 1.0 × 10□ CFU/mL of *P. aeruginosa*. The mixtures were subsequently incubated at 30°C and 180 r/min for 5 days. The control group of flasks containing only *P. aeruginosa* received the same treatment as described above. Cell densities of *P. aeruginosa* in the control and treatment groups were measured daily through counting CFU on TCBS agar plates from a series of ten-fold dilutions in sterile normal saline (Dong-ju, 2012). The bacteriolysis rate (%) is calculated according to the following formula (Zhang et al., 2022).

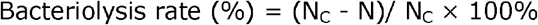

where N_C_ represents the cell density in the control group, and N indicates the cell densities in the treatment groups.

### 2.8 Bacterial pathogenicity and protective effect assay

#### 2.8.1 Experimental crabs

Healthy *E. sinensis* (22.12±0.6 g in weight) were provided by Nantong Duoruixian E-commerce Co., Ltd., Jiangsu, China. Following careful health assessment through physical observation and pathogen examination of a few individuals according to Ding et al. (2015) and Yang et al. (2018), the experimental crabs were acclimated in aerated tap water with pH7-8, dissolved-oxygen ≥ 6.0 mg/L, and total ammonia ≤ 0.2 mg/L at 28°C under a 12h light: 12h dark cycle for 14 days (Speights and McCoy, 2017; Xu et al., 2025).

#### 2.8.2 Bacterial pathogenicity assay

The pathogenicity of the isolate was examined by testing the median lethal dose (LD_50_) towards *E. sinensis* according to Cao et al. (2023). Prior to the LD_50_ test, the suspensions of the isolate were freshly prepared as described above (see section 2.6), and quantified as 5.0×10^4^, 5.0×10^5^, 5.0×10^6^, 5.0×10^7^, and 5.0×10^8^ CFU/mL through counting CFU on NA plates from a series of ten-fold dilutions in sterile normal saline. The LD_50_ test was carried out in 18 glass aquaria (60 cm × 40 cm × 40 cm; 10 crabs per aquarium) supplied with six tiles at the bottom and 10 L of aerated tap water at 28°C. These aquaria were randomly divided into one control and five treatment groups (three replicate aquaria per group). Crabs in the treatment groups were respectively intraperitoneally injected with 0.1 mL of 5.0×10^4^, 5.0×10^5^, 5.0×10^6^, 5.0×10^7^, and 5.0×10^8^ CFU/mL of the isolate suspension at the base of the third periopod. In contrast, the control group of crabs received intraperitoneal injection with 0.1 mL of sterile normal saline. During the test, crabs were fed using sterile-processed basal diet (with the formulation and proximate composition described previously (Cao et al., 2022), Hongxiang Feed Biotech. Co., Ltd., Jiangsu, China) at 8:00 and 18:00 up to apparent satiation at 3% total body weight (Fu et al., 2017), and kept at 28°C under a 12h light: 12h dark cycle for 7 days (Speights and McCoy, 2017). Residual food and feces as well as aquarium water exchange with fresh aerated tap water were carried out daily to ensure optimal survivals (Cheng et al., 2008). Dead crabs were immediately removed for bacterial isolation and identification according to Liu et al. (2024) to confirm if the mortality resulted from the isolate. Mortality of crabs in each group was recorded daily, and the LD_50_ value was estimated using the modified Kärber’s method (Zhang et al., 2022).

#### 2.8.3 Protection against P. aeruginosa infection in crabs assay

Prior to this assay, the suspensions of the isolate and *P. aeruginosa* strain HX-1 were freshly prepared as described above (see sections 2.2 and 2.6), and diluted to a final volume of 200 mL with distilled water to achieve 3.0×10^4^, 3.0×10^5^ and 3.0×10^6^ CFU/mL of the isolate and 2.5×10^7^ CFU/mL of *P. aeruginosa*. These dilutions were then manually added to 1.0 kg of the sterile-processed basal diet (Hongxiang Feed Biotech. Co., Ltd., Jiangsu, China), respectively, and the resulting mixtures were thoroughly blended in a drum mixer for 15 min, and subsequently air-dried under sterile conditions to achieve 6.0×10^3^, 6.0×10^4^, and 6.0×10^5^ CFU/g diet of the isolate-supplemented diet and 5.0×10^6^ CFU/g diet of *P. aeruginosa*-supplemented diet. The cell density in the feed was respectively quantified by counting CFU on NA plates from a series of ten-fold dilutions in sterile normal saline (Ariole et al., 2016). The assay was carried out in 12 glass aquaria (60 cm × 40 cm × 40 cm; 10 crabs per aquarium) supplied with three tiles at the bottom and 10 L of aerated tap water at 28°C. These aquaria were randomly divided into one control and three treatment groups (three replicate aquaria per group). Crabs in the treatment groups were respectively fed with 6.0×10^3^, 6.0×10^4^, and 6.0×10^5^ CFU/g diet of the isolate-supplemented diets. In contrast, the control group of crabs were fed with the sterile-processed basal diet. Following 20 days of feeding, crabs in each group were orally challenged with the *P. aeruginosa*-supplemented diet for 7 days. During the experiment, crabs were fed at 8:00 and 18:00 up to apparent satiation at 3% total body weight (Fu et al., 2017), and kept at 28°C under a 12h light: 12h dark cycle (Speights and McCoy, 2017). Residual food and feces as well as aquarium water exchange with fresh aerated tap water were carried out daily to assure optimal survivals (Cheng et al., 2008). Mortality of crabs in each group was recorded daily during the challenge period, and the relative percent survival (RPS) was calculated using the following formula (Souza et al., 2017).

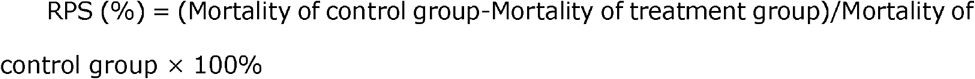

### 2.9 Statistical analysis

All data are presented as mean ± standard deviation, and analyzed using one-way analysis of variance (ANOVA) with Dunnett’s post hoc test in SPSS 19.0 software (SPSS, Inc., Chicago, IL, USA). Statistical significance is defined as *P* < 0.05.

## 3 Results

### 3.1 Bacterial isolation and identification

A facultatively predatory isolate, designated P2-4, was isolated from aquaculture sediment, and exhibited clear and visible plaques on the double-layer agar plate (Figure S1), similar to those showed by *bdellovibrios* (Cao et al., 2007). Isolate P2-4 grew well on nutrient agar, and were unable to grow on TCBS agar. Its partial 16S rRNA gene and whole-genome sequences were deposited in the GenBank database under accession nos. PV612026 and CM132631. Sequence analysis showed that isolate P2-4 shared a similarity of 97% to 99% with *M. varians* strains available in the GenBank database, and was further confirmed as a *M. varians* strain based on the phylogenetic analysis (Figure 1). The complete genome of isolate P2-4 consisted of one chromosome and one plasmid (Figure S2), with a whole genome size of 5,971,490 bp and an average GC content of 65.24% (Table 1). Totally, 75 tRNAs, 21 rRNAs, 34 sRNAs, 5238 CDS, 102 repeats, 1 prophage, 20 GI, and 23 IS were predicted in the genome of isolate P2-4 (Table 1). Furthermore, the ANI and dDDH values between isolate P2-4 and *M. varians* type strain CGMCC4.7419 were calculated as 95.93% and 79.50%, exceeding the species delineation thresholds of ANI (95%) and dDDH (70%) (Figure 2), further supporting isolate P2-4 as a *M. varians* strain.

**Table 1.** Genomic features of isolate P2-4.

| Feature | Genome |
| --- | --- |
| Genome size (bp) | 5,971,490 |
| Plasmid | 1 |
| Chromosome | 1 |
| GC average content (%) | 65.24 |
| Number of CDSs | 5238 |
| Gene average length (bp) | 1017.87 |
| Number of tRNAs | 75 |
| Number of 5S rRNAs | 7 |
| Number of 16S rRNAs | 7 |
| Number of 23S rRNAs | 7 |
| sRNAs | 34 |
| Repeated regions (%) | 0.23 |
| Number of repeats | 102 |
| Number of GI | 20 |
| Number of prophage | 1 |
| Number of IS | 23 |

GC, guanine–cytosine; CDS, coding sequences; GIs, genomic islands; IS, insertion sequence.

**Figure 1.**
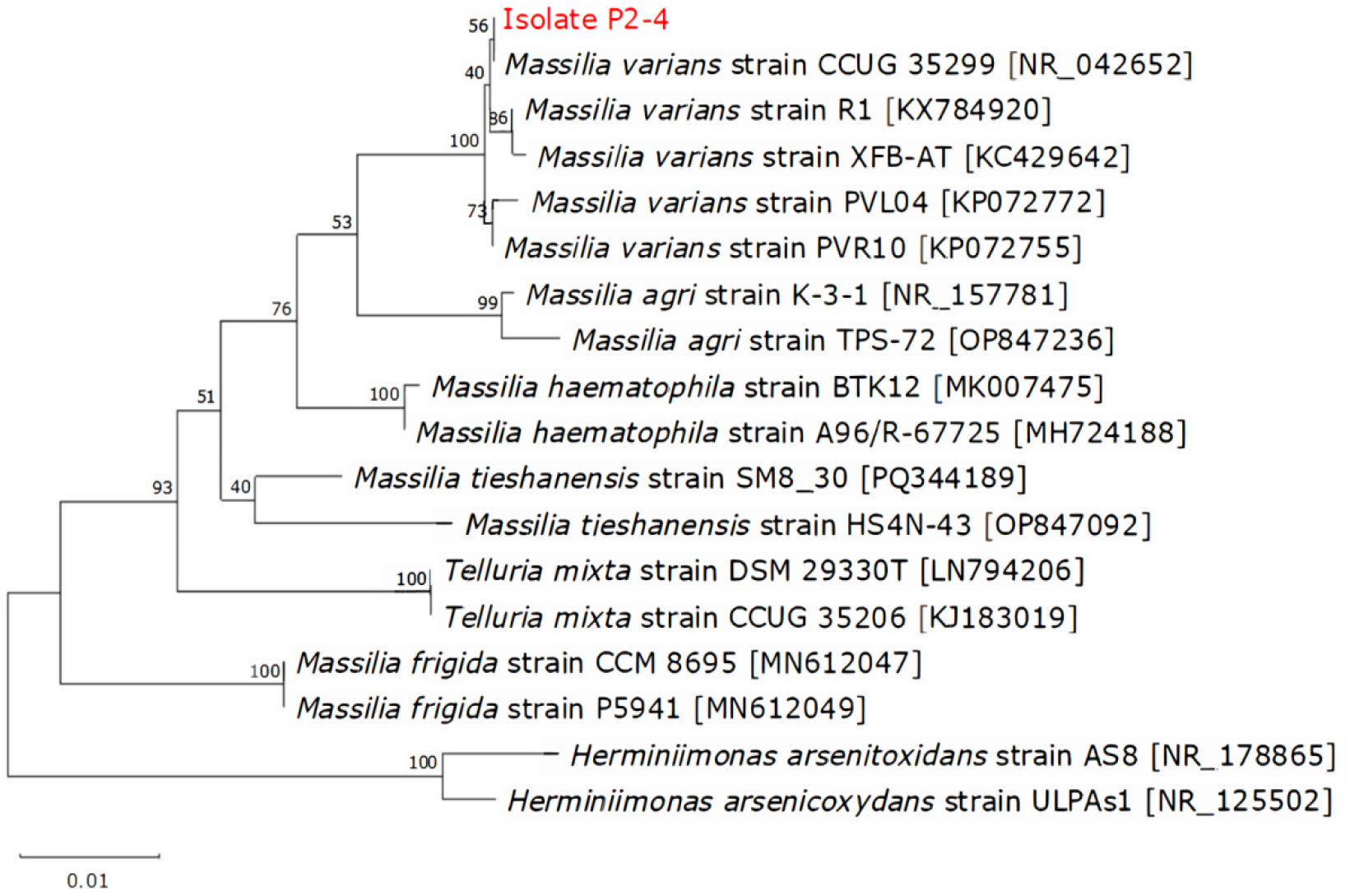
The 16S rRNA phylogenetic tree of isolate P2-4 and 17 known bacteria constructed using the neighbor-joining method. The bootstrap values (%) are shown beside the clades, accession numbers are indicated beside the names of strains, and scale bars represent distance values.

**Figure 2.**
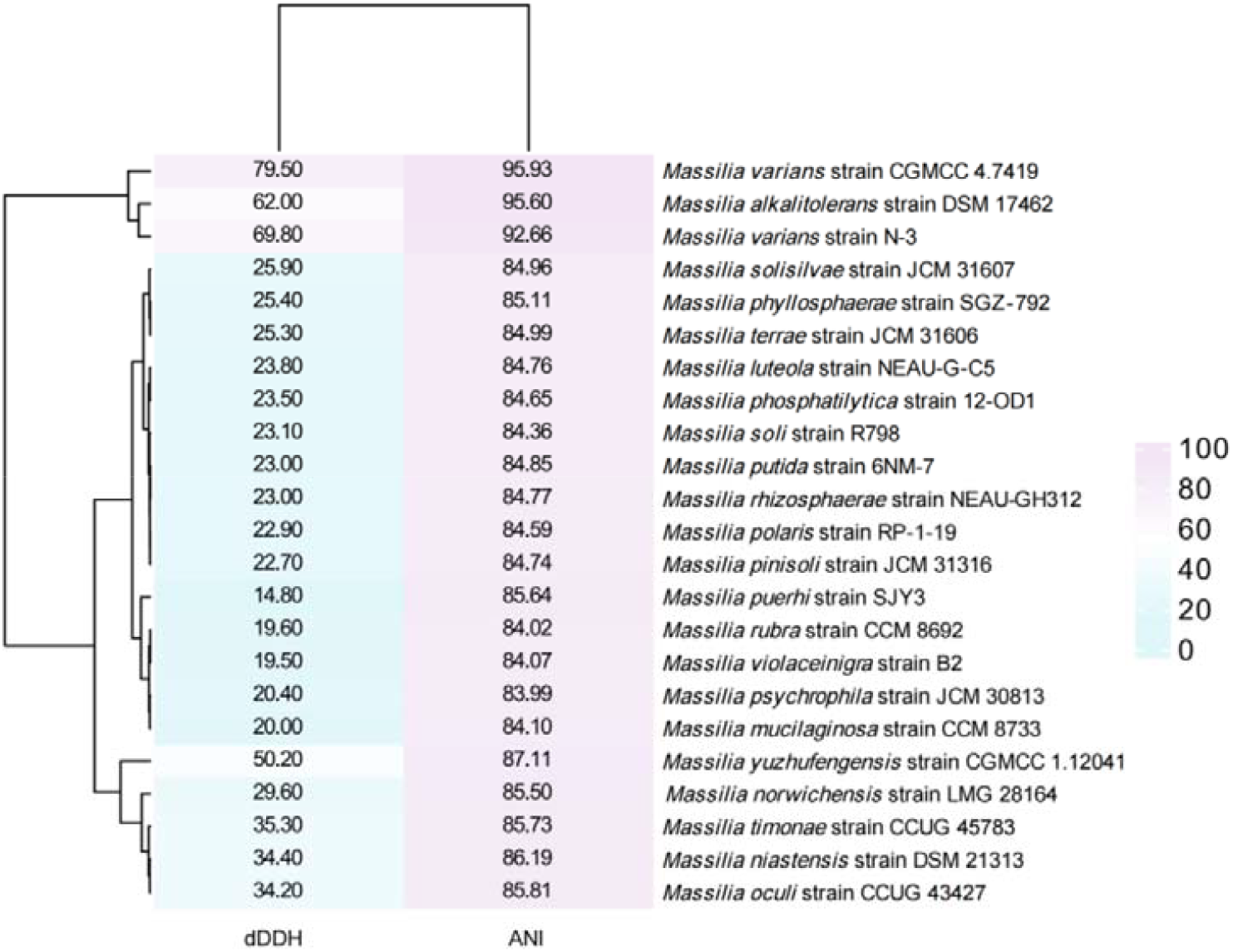
Heat map analysis of the genetic similarity between isolate P2-4 and other *Massilia* species based on the whole genome sequence. ANI, average nucleotide identity; dDDH, digital DNA– DNA hybridization.

Phenotypically, isolate P2-4 was Gram-negative rod-shaped, and formed round, slightly convex, smooth, moist, regular-edged, and pale-yellow colonies on the NA plate (Figure 3). It was positive for assimilation of glucose, maltose and malate, and was negative for nitrate reduction, indole production, glucose fermentation, arginine hydrolase, urease, β-glucosidase, and assimilation of arabinose, mannose, mannitol, N-acetyl-glucosamine, gluconate, capric acid, adipic acid, citrate, and phenylcetic acid (Table 2). These findings indicate that isolate P2-4 shows only 76.47% similarity to the reference strains of *M. varians* (Table 2), and differs from these reference strains in gelatinase production, glucose and arabinose fermentation, underscoring the inherent phenotypic diversity among *M. varians* strains.

**Table 2.** Phenotypic traits of isolate P2-4.

| Test | Reaction |  |
| --- | --- | --- |
|  | Isolate P2-4 | *M. varians*<sup>a</sup> |
| Nitrate reduction | - | - |
| Indole production | - | - |
| Glucose fermentation | - | - |
| Arginine hydrolase | - | ND |
| Urease | - | - |
| β-glucosidase | - | - |
| Gelatinase | - | + |
| β-galactosidase | - | - |
| Assimilation of |  |  |
| Glucose | + | - |
| Arabinose | - | + |
| Mannose | - | - |
| Mannitol | - | - |
| N-acetyl-D- (+)-glucosamine | - | ND |
| Maltose | + | + |
| Gluconate | - | - |
| Capric acid | - | ND |
| Adipic acid | - | - |
| Malate | + | ± |
| Citrate | - | - |
| Phenylacetic acid | - | + |

+, positive reaction; -, negative reaction; ±, positive or negative reactions; ND, not done. a, data previously reported (Cho et al., 2017; Chaudhary and Kim., 2017; Orthova et al., 2015; Weon et al., 2010).

**Figure 3.**
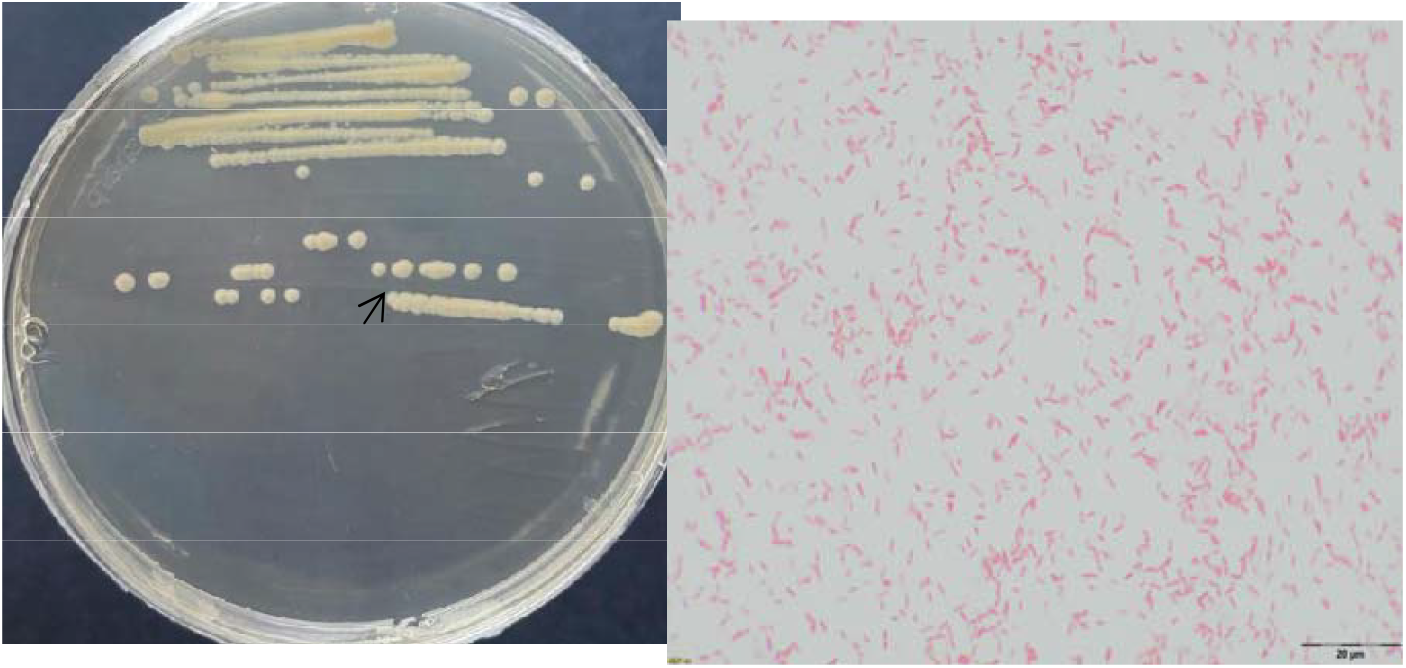
Morphological feature of isolate P2-4. (A) Colony morphology of isolate P2-4 cultured on a nutrient agar plate at 30 °C for 24 h. The arrow shows a round, slightly convex, smooth, moist, regular-edged, and pale yellow colony. (B) Microscopic morphology of isolate P2-4 (100×). The arrow shows a rod-shaped cell.

### 3.2 Bacterial virulence and bacteriolysis-related genes

Isolate P2-4 did not possess any virulence genes in VFDB, and harbored 19 bacteriolysis-related genes through genome blast against the COG and KEGG databases, including genes encoded pilus fibre forming protein PilA (*pilA*), pilus assembly ATPase PilB (*pilB*), pilin assembly protein PilC (*pilC*), prepilin peptidase PilD (*pilD*), prepilin leader peptidase PilF (*pilF*), signal transduction response regulator pilG (*pilG*), signal transduction response regulator PilH (*pilH*), pilus assembly protein PilM (*pilM*), pilus biogenesis protein PilO (*pilO*), pilus assembly protein PilP (*pilP*), pilus secretin PilQ (*pilQ*), pilus biogenesis protein PilS (*pilS*), pilus transcriptional regulator (*pilR*), pilus twitching motility protein PilT (*pilT*), lytic transglycosylase MltA (*mltA*), lytic transglycosylase MltB (*mltB*), lytic transglycosylase MltC (*mltC*), lytic transglycosylase MltD (*mltD)*, and endopeptidase DacB (*dacB*) that are crucial for invading and lysing prey cells (Table 3). These findings reveal the genetic basis for the nonpathogenicity and bacteriolysis of isolate P2-4.

**Table 3.** Potential bacteriolysis-related genes in isolate P2-4.

| Gene | Product | Identity (%) | Coverage (%) | E-value |
| --- | --- | --- | --- | --- |
| *pilA* | Pilus fibre forming protein PilA | 100 | 95.31 | 1.3e<sup>-31</sup> |
| *pilB* | Pilus assembly ATPase PilB | 99.7 | 99.83 | 0 |
| *pilC* | Pilin assembly protein PilC | 98.3 | 99.7 | 7.21e<sup>-276</sup> |
| *pilD* | Prepilin peptidase PilD | 99.7 | 98.97 | 7.02e<sup>-201</sup> |
| *pilF* | Prepilin leader peptidase PilF | 92.9 | 98.51 | 3.46e<sup>-165</sup> |
| *pilG* | Signal transduction<br>response regulator PilG | 100 | 99.16 | 6.95e<sup>-77</sup> |
| *pilH* | Signal transduction<br>response regulator PilH | 100 | 96.9 | 5.76e<sup>-80</sup> |
| *pilM* | Pilus assembly protein PilM | 99.7 | 99.72 | 1.01e<sup>-244</sup> |
| *pilO* | Pilus biogenesis protein PilO | 99.5 | 99.55 | 9.83e<sup>-149</sup> |
| *pilP* | Pilus assembly protein PilP | 98.4 | 99.45 | 2.41e<sup>-123</sup> |
| *pilQ* | Pilus secretin PilQ | 98.5 | 99.32 | 0 |
| *pilS* | Pilus biogenesis protein PilS | 98.2 | 99.82 | 0 |
| *pilR* | Pilus transcriptional regulator<br>PilR | 98.6 | 99.79 | 6.43e<sup>-312</sup> |
| *pilT* | Pilus twitching motility protein<br>PilT | 100 | 99.71 | 2.38e<sup>-244</sup> |
| *mltA* | Lytic transglycosylase MltA | 97.7 | 95.62 | 3.47e<sup>-256</sup> |
| *mltB* | Lytic transglycosylase MltB | 97.5 | 99.75 | 1.42e<sup>-265</sup> |
| *mltC* | Lytic transglycosylase MltC | 100 | 94.98 | 9.92e<sup>-189</sup> |
| *mltD* | Lytic transglycosylase MltD | 99.3 | 99.78 | 7.18e<sup>-308</sup> |
| *dacB* | Endopeptidase DacB | 96.7 | 99.19 | 0 |

### 3.3 Bacterial toxic metabolite production

Isolate P2-4 failed to produce toxic metabolites such as haemolysin, hydrogen sulfide, nitrite, and ammonia (Table 4). In addition, the control *R. azotoformans* strain SY5 was also unable to produce these toxic metabolites, consistent with its previous toxic metabolite production result (Cao et al., 2023), further indicating the reliability of toxic metabolite testing result with isolate P2-4. These findings indicate the non-toxic property of isolate P2-4.

**Table 4.** The toxic metabolites production of isolate P2-4.

| Toxic metabolite | Isolate P2-4 | *R. azotoformans* strain SY5 |
| --- | --- | --- |
| Haemolysin | - | - |
| Hydrogen sulfide | - | - |
| Nitrite | - | - |
| Ammonia | - | - |

-, negative reaction.

### 3.4 Bacterial bacteriolytic spectrum

Isolate P2-4 showed a wide bacteriolytic spectrum against *P. aeruginosa, S. algae, A. caviae, A. hydrophila*, and *P. damselae*. The maximum plaque number was recorded in *A. hydrophila* strain PX2-8 (385 PFU), significantly higher (*P* < 0.05) than that in *P. aeruginosa* strain HX-1 (315 PFU), *P. aeruginosa* strain FJ3-2 (270 PFU), *A. caviae* strain PX3-1 (268 PFU), *S. algae* strain RZ2-1 (244 PFU), *P. aeruginosa* strain FJ1-9 (225 PFU), *P. aeruginosa* strain PX3-3 (233 PFU), *S. algae* strain RZ2-2 (198 PFU), *A. caviae* strain PX2-5 (180 PFU), *P. damselae* strain YJ-1 (175 PFU), *P. aeruginosa* strain FJ1-5 (151 PFU), *S. algae* strain RZ3-4 (148 PFU), *P. aeruginosa* strain FJ1-2 (146 PFU), and *A. hydrophila* strain PX2-6 (121 PFU) (Figure 4). These findings reveal that isolate P2-4 possesses antibacterial potential against *P. aeruginosa, S. algae, A. caviae, A. hydrophila*, and *P. damselae*.

**Figure 4.**
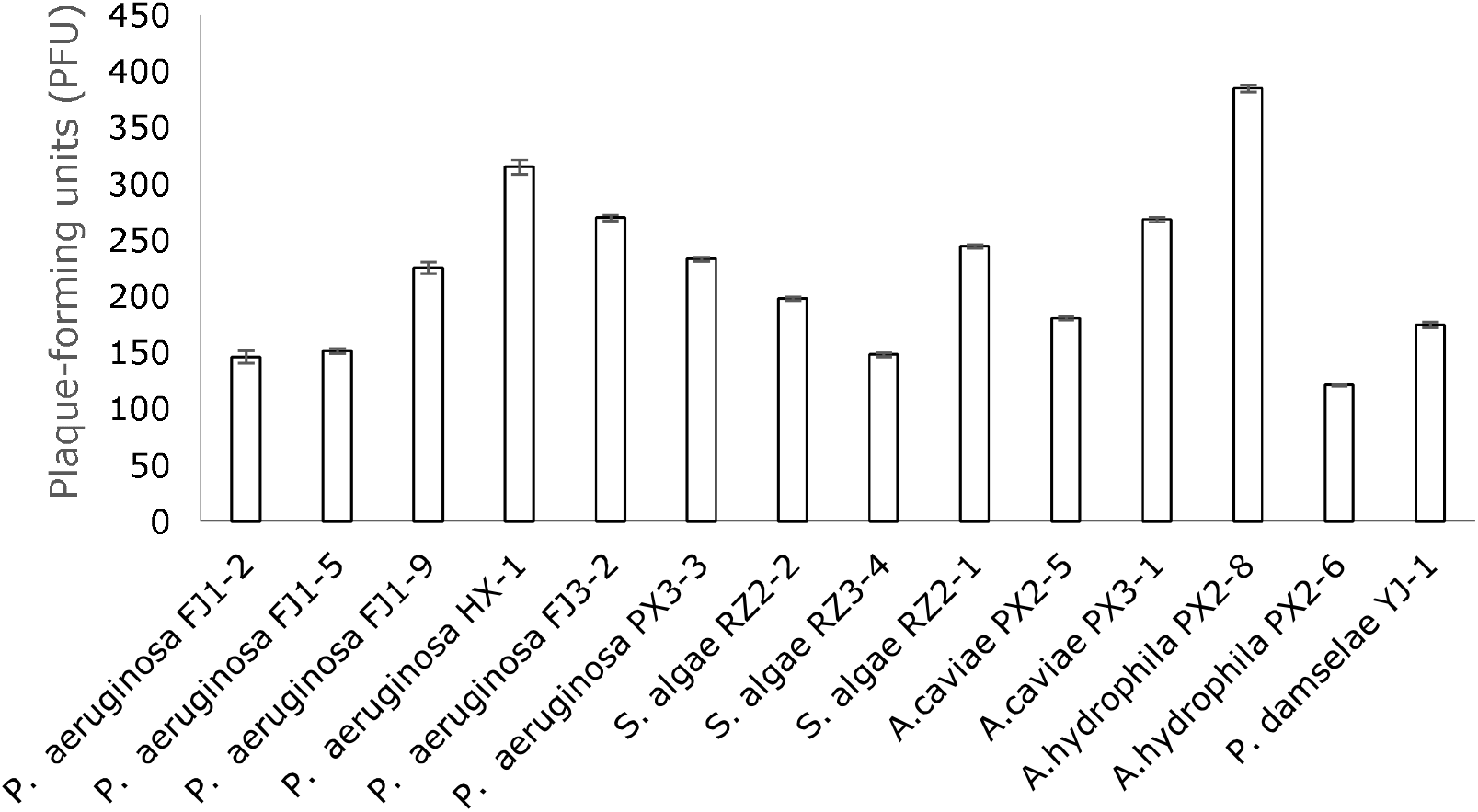
Bacteriolytic activity of isolate P2-4 against bacterial pathogens.

### 3.5 *In vitro* bacterial antibacterial effect

Compared to the control, the growth of *P. aeruginosa* treated with isolate P2-4 was significantly inhibited, and the cell density logarithms of *P. aeruginosa* were significantly reduced by 36.85% (*P* < 0.05), 50.89% (*P* < 0.05) and 68.17% (*P* < 0.05) following 5 days of co-culture with isolate P2-4 at 1.0×10^3^, 1.0×10□ and 1.0×10□ CFU/mL (Figure 5), obtaining bacteriolysis rates of 99.35%, 99.90% and 99.99% against *P. aeruginosa*. These findings demonstrate a significant antibacterial effect of isolate P2-4 against *P. aeruginosa*.

**Figure 5.**
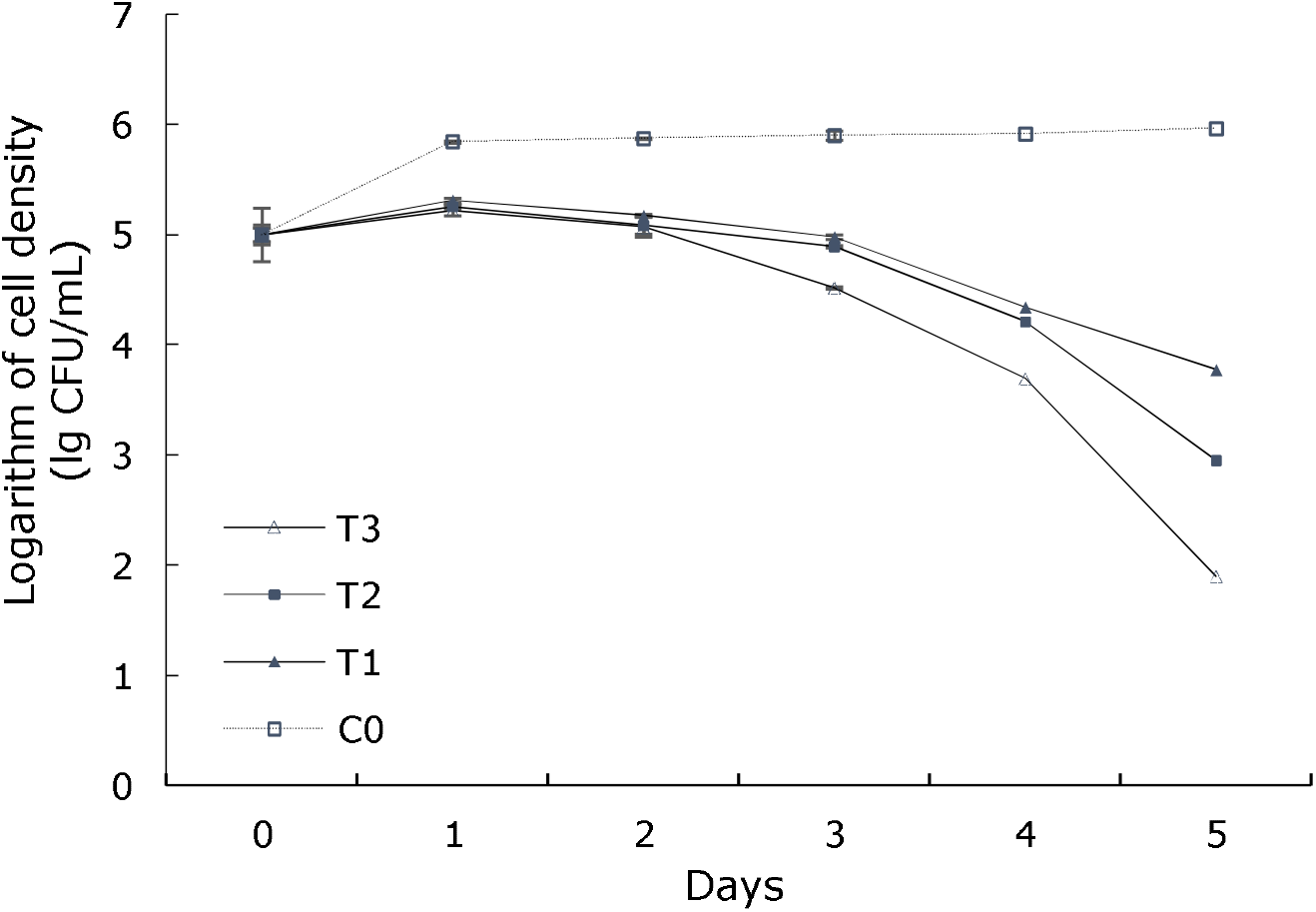
Effect of isolate P2-4 on the growth of *P. aeruginosa*. C0, *P. aeruginosa* only; T1, *P. aeruginosa +* 1.0×10^3^ CFU/mL of isolate P2-4; T2, *P. aeruginosa +* 1.0×10^4^ CFU/mL of isolate P2-4; T3, *P. aeruginosa +* 1.0×10^5^ CFU/mL of isolate P2-4.

### 3.6 Bacterial pathogenicity and protection against *P. aeruginosa* infection

No mortality or observable pathological signs were recorded in the control and 5.0×10^4^ ~ 5.0×10^8^ CFU/mL of isolate P2-4-treated crabs (data not shown), demonstrating the 7-day acute intraperitoneal LD_50_ value of isolate P2-4 as above 5.0 × 10^8^ CFU/mL in *E. sinensis*. These findings indicate isolate P2-4 as an avirulent strain towards *E. sinensis* based on the avirulent threshold (≥ 10^7^ CFU/mL) of acute LD_50_. Besides, crabs fed with isolate P2-4-supplemented diets exhibited significant improvement in survivals against *P. aeruginosa* challenge. Following 7 days of oral challenge with *P. aeruginosa*, mortality rates were significantly reduced by 36.67% (*P* < 0.05), 53.34% (*P* < 0.05) and 63.34% (*P* < 0.05) in crabs fed with 6.0×10^3^, 6.0×10□ and 6.0×10□ CFU/g diet of isolate P2-4-supplemented diets as compared to the control (Figure 6), obtaining the RPS values of isolate P2-4 at 6.0×10^3^, 6.0×10□ and 6.0×10□ CFU/g diet as 42.31%, 61.54% and 73.08%. In addition, the pathogenic *P. aeruginosa* could also be re-isolated from all dead crabs, which were confirmed through phenotypic and molecular identification (data not shown). These findings indicate that isolate P2-4 can confer significant protection of crabs against *P. aeruginosa* infection.

**Figure 6.**
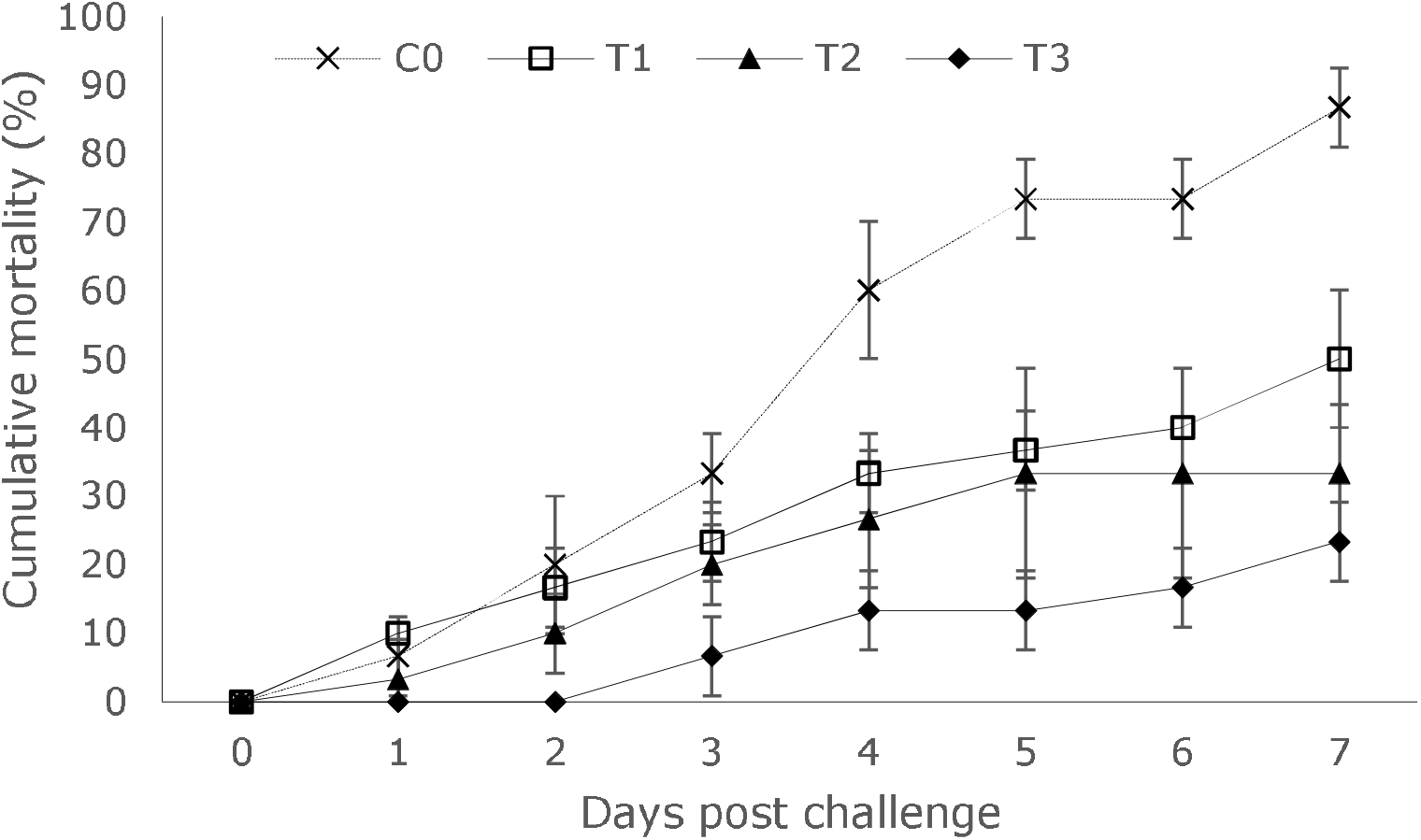
Protective effect of isolate P2-4 against *P. aeruginosa* infection in *E. sinensis*. C0, control crabs; T1, crabs fed with 6.0×10^3^ CFU/g diet of isolate P2-4-supplemented diets; T2, crabs fed with 6.0×10^4^ CFU/g diet of isolate P2-4-supplemented diets; T3, crabs fed with 6.0×10^5^ CFU/g diet of isolate P2-4-supplemented diets.

## 4 Discussion

Recently, environmental-friendly control agents have been proposed to control pathogenic *P. aeruginosa* in aquaculture, including the use of *Melaleuca alternifolia* essential oil, Yucca extract, *Chaetomorpha linum* extract, grape pomace flour, bacteriophages, *Bacillus coagulans, Padina boergesenii*, and levamisole (Koudou et al., 2020; Baldissera et al., 2019; El-Gammal et al., 2025; Ji et al., 2022; Naena, 2020; Ragunath and Ramasubramanian, 2022; Souza et al., 2017; Thanigaivel et al., 2023). Yet no reports are available on *M. varians* strains as biocontrol agents against pathogenic *P. aeruginosa* in aquaculture. In the present study, we identified a *M. varians* isolate P2-4 as a potential candidate for probiotic against crab-pathogenic *P. aeruginosa* through comprehensive analyses of genome-based bacteriolysis-related genes, safety assessment, as well as the *in vitro* and *in vivo* antibacterial effects. To our knowledge, this is the first report addressing a *M. varians* strain as a potential biocontrol agent against pathogenic *P. aeruginosa* in aquaculture.

In contrast to the obligate predator such as *Bdellovibrio* and like organisms (BALOs), facultative predators such as members of *Myxococcales, Streptomycetaceae*, and *Cytophagia* have a more widespread distribution in water ecosystems (Hungate et al., 2021), and can grow on nutrient agar (Seccareccia et al., 2015). Thus, we isolate facultative predatory bacteria from aquaculture sediment, and quantified isolate P2-4 using nutrient agar. However, isolate P2-4 showed no growth on TCBS agar. Thus, we used TCBS agar to quantify the cell density of *P. aeruginosa* that was co-cultured with isolate P2-4.

Given that 16S rRNA gene sequencing and phenotypic characterization analyses are insufficient for acute differentiation of *Massilia* species, whole genome sequencing is proposed for precise species identification (Baek et al., 2022). In the present study, whole genome-based analysis showed 95.93% ANI identity and 79.50% dDDH similarity to *M. varians* type strain CGMCC4.7419, taxonomically suggesting isolate P2-4 as a *M. varians* strain based on the species delineation thresholds of 95% ANI and 70% dDDH (Candeliere et al., 2021). However, isolate P2-4 exhibited only 76.47% phenotypic identity to *M. varians* reference strains, and differs from the clinical isolates of *M. varians* in arabinose and malate assimilation (Cho et al., 2017), revealing the lack of a large number of beneficial *M. varians* phenotypic data. Thus, there is an urgent need to collect extensive phenotypic trait data of beneficial *M. varians* to distinguish the difference between beneficial and potentially pathogenic strains of this species.

Recent studies have utilized whole genome sequencing to characterize the genomes of several *M. varians* strains. For example, *M. varians* strain CCUG 35299, isolated from the eye of a 90-year-old man in Tromso□, Norway (Kampfer et al., 2008), is composed of a chromosome with a whole genome size of 5,896,910 bp and a GC content of 65.5%, and possesses 67 tRNAs, 5 rRNAs, and 5,310 CDS (NCBI, 2024). *M. varians* strain N-3, isolated from Antarctic soil, consists of a single chromosome with a whole genome size of 5,575,375 bp and a GC content of 65.5%, and harbors 75 tRNAs, 21 rRNAs, and 4,941 CDS (NCBI, 2022). Yet no information is currently available regarding the genome of *M. varians* in aquaculture. In our study, the genome of isolate P2-4 from aquaculture sediment was firstly reported, and differed from *M. varians* strains CCUG 35299 and N-3 in the genomic features such as genome size and CDS number, revealing substantial genomic diversity among clinical and environmental isolates of *M. varians*. This is probably attributed to different environmental and nutritional stresses faced by these isolates (Pang et al., 2019).

Bacteriolysis-related genes are of great importance to predation in predatory bacteria (Oyedara et al., 2018). For example, type IV pili-encoding genes, such as *pilA, pilT, pilQ, pilM, pilS*, and *pilR*, are associated with adherence to and invasion of prey cells, and thus play a crucial role in predation of predatory bacteria (Evans et al., 2007; Mahmoud and Koval, 2010). The lytic transglycosylases and endopeptidases-encoding genes, such as *mltA, mltB, mltC, mltD*, and *dacB*, are involved in hydrolysing polysaccharide linkages of murein and attacking the peptide cross-links (Bernhardt and De Boer, 2004), resulting in relaxation of the peptidoglycanlayer and elimination of the prey cell septum obstruction (Dörr, 2023; Lerner et al., 2012). In the present study, besides *pilA, pilT, pilQ, pilM, pilS, pilR, mltA, mltB, mltC, mltD*, and *dacB* genes associated with predation and elimination of prey cells, other type IV pili-encoding genes such as *pilB, pilC, pilD, pilF, pilG, pilH, pilO*, and *pilP* were also present in isolate P2-4, probably contributing to the tight attachment and invasion of the predatory bacterial cells to prey cells (Mahmoud and Koval, 2010; Manetti and Spadafina, 2014). The presence of these bacteriolysis-related genes provides genetic explanation for the bacteriolysis of isolate P2-4.

Safety assessment of potential probiotic strains in aquaculture should be considered before their practical application (Verschuere et al., 2000). Gene mining using whole-genome data, coupled with experimental validation, is strongly recommended for probiotic safety assessment (Altavas et al., 2024; Wang et al., 2021). In our study, isolate P2-4 did not possess any known virulence genes through genome blast against VFDB database, was unable to produce toxic haemolysin, hydrogen sulfide, nitrite, and ammonia that could cause systemic damage and toxic effects on crabs (Abd-El-Malek, 2017; Koo et al., 2005; Xu et al., 2014), suggesting that isolate P2-4 has no pathogenic potential. In addition, the acute LD_50_ value is regarded as a precise indicator for safety assessment of probiotics in aquaculture (Turnip et al., 2018; Zhou et al., 2000). Bacterial isolates are regarded as avirulent based on LD_50_ values of above 10^7.0^ CFU/mL (Triyaningsih et al., 2014). Isolate P2-4 showed an acute intraperitoneal LD_50_ value of above 5.0 × 10^8^ CFU/mL in *E. sinensis*, further supporting it as an avirulent strain. However, it should be noted that *M. varians* isolates have been occasionally isolated from clinical specimens (eye and finger) in humans (Cho et al., 2017; Kampfer et al., 2008), but no evidence suggests the pathogenicity of these isolates in humans and nonhuman animals. Similar issues also exist in *Enterococcus faecalis* and *Weissella confusa* that have been proposed as probiotics, but also opportunistically causes infections in humans or animals (De Almeida et al., 2018; Fairfax et al., 2014). Thus, probiotic strains within the same species are still promising candidate for the control of infectious agents and treatment of disease (Chauhan and Singh, 2019).

Obligate predatory bacteria are considered to be economically rewarding as they can prey upon and kill a broad spectrum of multiple bacterial pathogens (Damron and Barbier, 2013; Melander et al., 2018). For instance, *Bdellovibrio* sp. isolate T1 displays predation against 45.45% of the 11 test prey strains (Ezzedine et al., 2020). *Bdellovibrio* sp. isolate YBD-1 shows bacteriolysis against 92.31% of the 13 test prey strains (Xi et al., 2024). In the present study, isolate P2-4 exhibited bacteriolytic activities against all test *P. aeruginosa, A. caviae, A. hydrophila, P. damselae*, and *S. algae* pathogens that pose a significant threat to aquaculture (Dehkordi et al., 2022; Lattos et al., 2022; Tao et al., 2026; Zago, 2020), suggesting a wider bacteriolytic spectrum than the obligate *Bdellovibrio* sp. isolates T1 and YBD-1 (Ezzedine et al., 2020; Xi et al., 2024). The wide bacteriolytic spectrum demonstrated by isolate P2-4 suggests its potential as a candidate for probiotics to inhibit *P. aeruginosa, A. caviae, A. hydrophila, P. damselae*, and *S. algae* pathogens in aquaculture.

*P. aeruginosa* is an emerging but underestimated bacterial pathogen in *E. sinensis* aquaculture (He et al., 2026). To assess the practical application value of isolate P2-4, our study examined its antibacterial efficacy *in vitro* and *in vivo* against the *E. sinensis*-pathogenic *P. aeruginosa*. The findings indicated that isolate P2-4 at 1.0×10^3^ to 1.0×10□ CFU/mL displayed bacteriolysis rates of 99.35% to 99.99% against the crab-pathogenic *P. aeruginosa*. This is probably due to the widespread presence of *pil* genes encoding type IV pili that promote the isolate’s strong predatory activity in liquid prey cultures (Evans et al., 2007). Furthermore, isolate P2-4 at 6.0×10^3^ to 6.0×10^5^ CFU/g diet also led to RPS of 42.31% to 73.08% against *P. aeruginosa* infection in *E. sinensis*. This can be presumptively attributed to reduction of the pathogen numbers due to bacteriolysis by isolate P2-4 (Cao et al., 2014a).

## 5. Conclusion

The present study for the first time reveals the *M. varians* isolate (P2-4) as a potential biocontrol agent against *P. aeruginosa* in aquaculture. The absence of virulence genes, a large presence of bacteriolysis-related genes, nonpathogenicity and the *in vitro* and *in vivo* significant antibacterial efficacy in isolate P2-4 support this bacterium as a potential candidate for probiotic against pathogenic *P. aeruginosa* in aquaculture.

## Data availability statement

The datasets presented in this study can be found in online repositories. The names of the repository/repositories and accession number(s) can be found in the article/Supplementary material. Requests to access the datasets should be directed to the corresponding author.

## Author contribution

LY: writing the original draft. YY: investigation, data curation, formal analysis. LM: writing-review and editing. CS: data curation, formal analysis. CH: writing the original draft, project administration. GC: resources, funding acquisition. YW: formal analysis, funding acquisition.

## Funding

This work was supported by the Earmarked Fund for China Agriculture Research System (No. CARS-48).

## Conflict of Interest

The authors declare no conflicts of interest.

## Generative AI statement

The author(s) declared that Generative AI was not used in the creation of this manuscript. Any alternative text (alt text) provided alongside figures in this article has been generated by Frontiers with the support of artificial intelligence and reasonable efforts have been made to ensure accuracy, including review by the authors wherever possible. If you identify any issues, please contact us.

## Publisher’s note

All claims expressed in this article are solely those of the authors and do not necessarily represent those of their affiliated organizations, or those of the publisher, the editors and the reviewers. Any product that may be evaluated in this article, or claim that may be made by its manufacturer, is not guaranteed or endorsed by the publisher.

## Supplementary material

The Supplementary material for this article can be found online in the Supplementary material section.

Figure S1. The plaques of isolate P2-4 formed on the double-layer agar plate after 5 days of incubation at 30°C, using *P. aeruginosa* as the prey bacterium.

Figure S2. Circular complete genome map of isolate P2-4.

## Notes

### Competing Interest Statement

The authors have declared no competing interest.

## Reference

Abd-El-Malek, A. M. (2017). Incidence and virulence characteristics of Aeromonas spp. in fish. Vet. World. 10(1), 34–37. doi: 10.14202/vetworld.2017.34-37

Agbekpornu, H., Yuan, X., Ming, J., and Zhu, W. (2018). Marketing of freshwater Chinese mitten crab and support to the mitten crab farmers in China. Asian J. Agric. Ext. Econ. Sociol. 27(3), 1–10. doi: 10.9734/AJAEES/2018/44041

Ali, N. G., Ali, T. E. S., Aboyadak, I. M., and Elbakry, M. A. (2021). Controlling Pseudomonas aeruginosa infection in Oreochromis niloticus spawners by cefotaxime sodium. Aquaculture 544(1), 737107. doi: 10.1016/j.aquaculture.2021.737107

Altavas, P. J. d.R., Amoranto, M. B. C., Kim, S. H., Kang, D. K., Balolong, M. P., and Dalmacio, L. M. M. (2024). Safety assessment of five candidate probiotic lactobacilli using comparative genome analysis. Access Microbiol. 6(1). doi: 10.1099/acmi.0.000715.v4

Anokyewaa, M. A., Amoah, K., Li, Y., Lu, Y., Kuebutornye, F. K. A., Asiedu, B., et al. (2021). Prevalence of virulence genes and antibiotic susceptibility of Bacillus used in commercial aquaculture probiotics in China. Aquac. Rep. 21(3), 100784. doi: 10.1016/j.aqrep.2021.100784

Ariole, C. N., and Eddo, T. T. (2016). Effect of an indigenous probiotic (Shewanella algae) isolated from healthy shrimp (Penaeus monodon) intestine on Clarias gariepinus. Aquacult Int, 5 (36).,1–9. doi: 10.5376/ija.2015.05.0036

Baek, J. H., Baek, W., Ruan, W., Jung, H. S., Lee, S. C., and Jeon, C. O. (2022). Massilia soli sp. nov., isolated from soil. Int. J. Syst. Evol. Microbiol. 72(2). doi: 10.1099/ijsem.0.005227

Baldissera, M. D., Souza, C. F., Descovi, S. N., Verdi, C. M., Zeppenfeld, C. C., De Lima Silva, L., et al. (2019). Effects of dietary grape pomace flour on the purinergic signaling and inflammatory response of grass carp experimentally infected with Pseudomonas aeruginosa. Aquaculture 503, 217–224. doi: 10.1016/j.aquaculture.2019.01.015

Bauer, A., and Forchhammer, K. (2021). Bacterial predation on Cyanobacteria. Microb. Physiol. 31(2), 99–108. doi: 10.1159/000516427

Benson, G. (1999). Tandem repeats finder: a program to analyze DNA sequences. Nucleic Acids Res. 27(2), 573–580. doi: 10.1093/nar/27.2.573

Bernhardt, T. G., and De Boer, P. A. J. (2004). Screening for synthetic lethal mutants in Escherichia coli and identification of EnvC (YibP) as a periplasmic septal ring factor with murein hydrolase activity. Mol. Microbiol. 52(5), 1255–1269. doi: 10.1111/j.1365-2958.2004.04063.x

Besemer, J., and Borodovsky, M. (2005). GeneMark: web software for gene finding in prokaryotes, eukaryotes and viruses. Nucleic Acids Res. 33, W451–W454. doi: 10.1093/nar/gki487

Candeliere, F., Raimondi, S., Spampinato, G., Tay, M. Y. F., Amaretti, A., Schlundt, J., et al. (2021). Comparative genomics of Leuconostoc carnosum. Front. Microbiol. 11, 605127. doi: 10.3389/fmicb.2020.605127

Cao, H., An, J., Zheng, W., and He, S. (2015). Vibrio cholerae pathogen from the freshwater-cultured whiteleg shrimp Penaeus vannamei and control with Bdellovibrio bacteriovorus. J. Invertebr. Pathol. 130, 13–20. doi: 10.1016/j.jip.2015.06.002

Cao, H., He, S., Lu, L., Yang, X., and Chen, B. (2014a). Identification of a Proteus penneri isolate as the causal agent of red body disease of the cultured white shrimp Penaeus vannamei and its control with Bdellovibrio bacteriovorus. Anton. Leeuw. 105, 423–430. doi: 10.1007/s10482-013-0079-y

Cao, H., He, S., Wang, H., Hou, S., Lu, L., and Yang, X. (2012). Bdellovibrios, potential biocontrol bacteria against pathogenic Aeromonas hydrophila. Vet. Microbiol. 154 (3-4), 413–418. doi: 10.1016/j.vetmic.2011.07.032

Cao, H., Hou, S., He, S., Lu, L., and Yang, X. (2014b). Identification of a Bacteriovorax sp. isolate as a potential biocontrol bacterium against snakehead fish-pathogenic Aeromonas veronii. J. Fish Dis. 37 (3), 283–289. doi: 10.1111/jfd.12120

Cao, H., Wang, H., Yu, J., An, J., and Chen, J. (2019). Encapsulated Bdellovibrio powder as a potential bio-disinfectant against whiteleg shrimp-pathogenic vibrios. Microorganisms 7 (8), 244. doi: 10.3390/microorganisms7080244

Cao, H., Yang, X., Qian, Y., and Deng, L. (2007). Isolation of Bdellovibrio bacteria from the gut of Carassius auratus gibelio and the study of its biological characteristics. Microbiol. China 34, 52–56.

Cao, H., Zhang, S., An, J., Diao, J., Xu, L., and Gai, C. (2022). Rhodobacter azotoformans supplementation improves defense ability of Chinese mitten crab Eriocheir sinensis against citrobacteriosis. Fish Shellfish Immunol. 131, 991–998. doi: 10.1016/j.fsi.2022.11.012

Cao, H., Zhang, S., Zheng, X., Xu, L., Diao, J., Wang, Y., et al. (2023). Safety assessment of Rhodobacter azotoformans SY5 for potential application in Chinese mitten crab Eriocheir sinensis. Benef. Microbes 14(3), 641–651. doi: 10.1163/18762891-20230086

Castro-Escarpulli, G., Figueras, M. J., Aguilera-Arreola, G., Soler, L., Fernández-Rendón, E., Aparicio, G. O., et al. (2003). Characterisation of Aeromonas spp. isolated from frozen fish intended for human consumption in Mexico. Int. J. Food Microbiol. 84 (1), 41–49. doi: 10.1016/S0168-1605(02)00393-8

Chan, P. P., and Lowe, T. M. (2019). tRNAscan-SE: Searching for tRNA genes in genomic sequences. Methods Mol. Biol. 1962, 1–14. doi: 10.1007/978-1-4939-9173-0_1

Chaudhary, D. K., and Kim, J. (2017). Massilia agri sp. nov., isolated from reclaimed grassland soil. Int. J. Syst. Evol. Microbiol. 67(8), 2696–2703. doi: 10.1099/ijsem.0.002002

Chauhan, A., and Singh, R. (2019). Probiotics in aquaculture: a promising emerging alternative approach. Symbiosis 77, 99–113. doi: 10.1007/s13199-018-0580-1

Cheng, Y., Wu, X., Yang, X., and Hines, A. H. (2008). Current trends in hatchery techniques and stock enhancement for Chinese mitten crab, Eriocheir japonica sinensis. Rev. Fish. Sci. 16 (1-3), 377–384. doi: 10.1080/10641260701681698

Cho, J., Kim, K. H., Kim, J. O., Hong, J. S., Jeong, S. H., and Lee, K. (2017). Massilia varians isolated from a clinical specimen. Infect. Chemother. 49 (3), 219–222. doi: 10.3947/ic.2017.49.3.219

Damron, F. H., and Barbier, M. (2013). Predatory bacteria: living antibiotics, biocontrol agents, or probiotics? Postdoc J. 1(12), 1–15. doi: 10.14304/SURYA.JPR.V1N12.2

De Almeida, C. V., Taddei, A., and Amedei, A. (2018). The controversial role of Enterococcus faecalis in colorectal cancer. Ther. Adv. Gastroenterol. 11, 1756284818783606. doi: 10.1177/1756284818783606

Dehkordi, S. M. H., Anvar, S. A., Rahimi, E., Ahari, H., and Ataee, M. (2022). Prevalence, phenotypic and genotypic diversity, antibiotic resistance, and frequency of virulence genes in Pseudomonas aeruginosa isolated from shrimps. Aquac. Int. 30, 131–156. doi: 10.1007/s10499-021-00798-z

Devakumar, D., Jayanthi, J., and Ragunathan, M. (2013). Herbal alternate to Pseudomonas aeruginosa infection in a freshwater crab, Oziotelphusa senex senex. Aquaculture 2, 2142–2147.

Ding, Z., Yao, Y., Zhang, F., Wan, J., Sun, M., Liu, H., et al. (2015). The first detection of white spot syndrome virus in naturally infected cultured Chinese mitten crabs, Eriocheir sinensis in China. J. Virol. Methods 220, 49–54. doi: 10.1016/j.jviromet.2015.04.011

Dong-ju Kim, (2012). Relation of microbial biomass to counting units for Pseudomonas aeruginosa. Afr. J. Microbiol. Res. 6 (21), 4620–4622. doi: 10.5897/AJMR10.902

Dörr, T. (2023). Cleave a septum, leave a cell: Bdellovibrio bacteriovorus secretes a specialized lytic transglycosylase to clear prey cell septum obstruction. J. Bacteriol. 205 (4), e0007423. doi: 10.1128/jb.00074-23

Eifrig, C. W. G., Scott, I. U., Flynn, H. W., and Miller, D. (2003). Endophthalmitis caused by Pseudomonas aeruginosa. Ophthalmology 110 (9), 1714–1717. doi: 10.1016/S0161-6420(03)00572-4

El-Gammal, G. A., El-Gamal, A. M., Rashed, M. A., Kassab, A. S., Saif, A. S., and Fadl, S. E. (2025). An experimental study of levamisole incorporated diet on fish health and resistance against Pseudomonas aeruginosa isolated from Oreochromas niloticus. Sci. Rep. 15 (1), 14658. doi: 10.1038/s41598-025-96914-7

El-Tarabili, R. M., Eid, H. M., Elghayaty, H. A. A., and Zaghloul, E. M. (2023). Detection of pelA and associated virulence genes in emerging multidrug-and extensively drug-resistant (MDR and XDR) Pseudomonas aeruginosa isolated from Oreochromis niloticus. Bulg. J. Vet. Med. 26 (4), 524–541. doi: 10.15547/bjvm.2021-0061

Evans, K. J., Lambert, C., and Sockett, R. E. (2007). Predation by Bdellovibrio bacteriovorus HD100 requires type IV pili. J. Bacteriol. 189 (13), 4850–4859. doi: 10.1128/JB.01942-06

Ezzedine, J. A., Pavard, G., Gardillon, M., and Jacquet, S. (2020). Bdellovibrio sp: An important bacterial predator in lake geneva? J. Microbiol. Biotechnol. 5(1), 000157. doi:10.23880/oajmb-16000157

Fairfax, M. R., Lephart, P. R., and Salimnia, H. (2014). Weissella confusa: problems with identification of an opportunistic pathogen that has been found in fermented foods and proposed as a probiotic. Front. Microbiol. 5. doi: 10.3389/fmicb.2014.00254

Fong, I. W., and Tomkins, K. B. (1985). Review of Pseudomonas aeruginosa meningitis with special emphasis on treatment with ceftazidime. Clin. Infect. Dis. 7 (5), 604–612. doi: 10.1093/clinids/7.5.604

Fouts, D. E. (2006). Phage_Finder: Automated identification and classification of prophage regions in complete bacterial genome sequences. Nucleic Acids Res. 34 (20), 5839–5851. doi: 10.1093/nar/gkl732

Frascaroli, G., Roberts, J., Hunter, C., and Escudero, A. (2024). Removal efficiencies of seven frequently detected antibiotics and related physiological responses in three microalgae species. Environ Sci Pollut R, 31(9), 14178–14190. doi: 10.1007/s11356-024-32026-5

Fu, L., Zhou, G., Pan, J., Li, Y., Lu, Q., Zhou, J., et al. (2017). Effects of Astragalus polysaccharides on antioxidant abilities and non-specific immune responses of Chinese mitten crab, Eriocheir sinensis. Aquac. Int. 25, 1333–1343. doi: 10.1007/s10499-017-0117-2

Gai, C., Zheng, X., Xu, L., Diao, J., Ye, H., Cao, H., and Yu, X. (2023). Haemolytic Bacillus subtilis, a potential pathogen of hemorrhage disease in farmed yellow catfish Pelteobagrus fulvidraco. Isr. J. Aquacult. Bamidgeh, 75(2), 1–12. doi: 10.46989/001c.84493

He, L., Yang, Y., Liu, Y., Liu, M., Cao, H., Gai, C., et al. (2026). Pseudomonas aeruginosa in Chinese mitten crab Eriocheir sinensis: Identification, pathogenic potential and control with Bdellovibrio powder. J. Invertebr. Pathol. 217, 108604. doi: 10.1016/j.jip.2026.108604

Huang, X., Gu, Y., Zhou, H., Xu, L., Cao, H., and Gai, C. (2020). Acinetobacter venetianus, a potential pathogen of red leg disease in freshwater-cultured whiteleg shrimp Penaeus vannamei. Aquac. Rep. 18, 100543. doi: 10.1016/j.aqrep.2020.100543

Hungate, B. A., Marks, J. C., Power, M. E., Schwartz, E., Van Groenigen, K. J., Blazewicz, S. J., et al. (2021). The functional significance of bacterial predators. mBio 12 (2), e00466–21. doi: 10.1128/mBio.00466-21

Ji, T., Cao, Y., Cao, Q., Zhang, Y., and Yang, H. (2022). The antagonistic effect and protective efficacy of gram-positive probiotics Bacillus coagulans to newly identified pathogens Pseudomonas aeruginosa in crucian carp Carassius auratus gibelio. Aquac. Rep. 24, 101126. doi: 10.1016/j.aqrep.2022.101126

Kamada, S., Wakabayashi, R., and Naganuma, T. (2023). Phylogenetic revisit to a review on predatory bacteria. Microorganisms 11 (7), 1673. doi: 10.3390/microorganisms11071673

Kampfer, P., Falsen, E., and Busse, H.-J. (2008). Naxibacter varians sp. nov. and Naxibacter haematophilus sp. nov., and emended description of the genus Naxibacter. Int. J. Syst. Evol. Microbiol. 58 (7), 1680–1684. doi: 10.1099/ijs.0.65516-0

Kanehisa, M. (2000). KEGG: Kyoto encyclopedia of genes and genomes. Nucleic Acids Res. 28, 27–30. doi: 10.1093/nar/28.1.27

Kanehisa, M., and Goto, S. (2000). KEGG: kyoto encyclopedia of genes and genomes. Nucleic Acids Res., 28(1), 27–30. doi: 10.1093/nar/28.1.27

Kang, M. S., Yeu, J. E., and Hong, S. P. (2019). Safety evaluation of oral care probiotics Weissella cibaria CMU and CMS1 by phenotypic and genotypic analysis. Int. J. Mol. Sci. 20 (11), 2693. doi: 10.3390/ijms20112693

Kholil, Md. I., Hossain, Md. M. M., Neowajh, Md. S., Islam, Md. S., and Kabir, M. (2015). Comparative efficiency of some commercial antibiotics against Pseudomonas infection in fish. Int. J. Fish. Aquat. Stud. 2 (3), 114–117. doi: 10.22271/fish.2015.v2.i3b.225

Koo, J. G., Kim, S. G., Jee, J. H., Kim, J. M., Bai, S. C., and Kang, J. C. (2005). Effects of ammonia and nitrite on survival, growth and moulting in juvenile tiger crab, Orithyia sinica (Linnaeus). Aquac. Res. 36 (1), 79–85. doi: 10.1111/j.1365-2109.2004.01187.x

Koren, S., Walenz, B. P., Berlin, K., Miller, J. R., Bergman, N. H., and Phillippy, A. M. (2017). Canu: scalable and accurate long-read assembly via adaptive k-mer weighting and repeat separation. Genome Res. 27, 722–736. doi: 10.1101/gr.215087.116

Koudou, A. A., Kakou-Ngazoa, S. E,, Addablah, A., Allali, K. B., Aoussi, S., Diallo, A. H., et al. (2020). Biocontrôle de l’infection à Pseudomonas aeruginosa multi-résistant par les bactériophages en aquaculture en Côte d’Ivoire. J. Appl. Biosci. 154, 15940–15949. doi: 10.35759/JABs.154.10

Krzywinski, M., Schein, J., Birol, İ., Connors, J., Gascoyne, R., Horsman, D., et al. (2009). Circos: An information aesthetic for comparative genomics. Genome Res. 19, 1639–1645. doi: 10.1101/gr.092759.109

Lattos, A., Giantsis, I. A., Tsavea, E., Kolygas, M., Athanassopoulou, F., and Bitchava, K. (2022). Virulence genes and in vitro antibiotic profile of Photobacterium damselae strains, isolated from fish reared in greek aquaculture facilities. Animals 12 (22), 3133. doi: 10.3390/ani12223133

Lerner, T. R., Lovering, A. L., Bui, N. K., Uchida, K., Aizawa, S. I., Vollmer, W., et al. (2012). Specialized peptidoglycan hydrolases sculpt the intra-bacterial niche of predatory Bdellovibrio and increase population fitness. PLoS Pathog. 8 (2), e1002524. doi: 10.1371/journal.ppat.1002524

Li, C., Cao, P., Du, C., Zhang, X., Bing, H., Li, L., et al. (2021). Massilia rhizosphaerae sp. nov., a rice-associated rhizobacterium with antibacterial activity against Ralstonia solanacearum. Int. J. Syst. Evol. Microbiol. 71 (9). doi: 10.1099/ijsem.0.005009

Li, K., Liu, X., Zuo, W., and Wu, N. (2023). Whole-genome sequencing of a multidrug-resistant strain: delftia acidovorans B408. Biochem. Genet. 61 (3), 1086–1096. doi: 10.1007/s10528-022-10306-4

Liao, C., Huang, X., Wang, Q., Yao, D., and Lu, W. (2022). Virulence factors of Pseudomonas aeruginosa and antivirulence strategies to combat its drug resistance. Front. Cell. Infect. Microbiol. 12, 926758. doi: 10.3389/fcimb.2022.926758

Lindquist, D., Murrill, D., Burran, W. P., Winans, G., Janda, J. M., and Probert, W. (2003). Characteristics of Massilia timonae and Massilia timonae-like isolates from human patients, with an emended description of the species. J. Clin. Microbiol., 41(1), 192–196. doi: 10.1128/JCM.41.1.192-196.2003

Liu, B., Zheng, D., Jin, Q., Chen, L., and Yang, J. (2019). VFDB 2019: a comparative pathogenomic platform with an interactive web interface. Nucleic Acids Res. 47 (D1), D687–D692. doi: 10.1093/nar/gky1080

Liu, J., Li, J., Huang, X., Chen, B., An, J., and Cao, H. (2022). Effect of dietary Bdellovibrio powder on non-specific immunity, antioxidant ability and disease resistance of Chinese mitten crab Eriocheir sinensis. Iran. J. Fish. Sci. 21, 948–965.

Liu, J., Teng, C., Zheng, X., Xu, L., Cao, H., and Gai, C. (2024). Shewanella putrefaciens as an emerging pathogen of hepatopancreas necrosis disease in Chinese mitten crab Eriocheir sinensis. Dis. Aquat. Organ. 159, 143–152. doi: 10.3354/dao03811

Luo, R., Liu, B., Xie, Y., Li, Z., Huang, W., Yuan, J., et al. (2012). SOAPdenovo2: an empirically improved memory-efficient short-read de novo assembler. Gigascience 1 (1), 2047-217X-1–18. doi:10.1186/2047-217X-1-18

Mahmoud, K. K., and Koval, S. F. (2010). Characterization of type IV pili in the life cycle of the predator bacterium Bdellovibrio. Microbiology 156, 1040–1051. doi:10.1099/mic.0.036137-0

Manetti, A. G. O., and Spadafina, T. (2014). The role of pili in the formation of biofilm and bacterial communities. In: Bacterial pili: structure, synthesis and role in disease, eds. M. A. Barocchi and J. L. Telford (Wallingford: CABI), 151–164. doi:10.1079/9781780642550.0151

Melander, R. J., Zurawski, D. V., and Melander, C. (2018). Narrow-spectrum antibacterial agents. MedChemComm 9 (1), 12–21. doi:10.1039/C7MD00528H

Mohsenipour, Z., Arazi, P., Skurnik, M., Jahanbin, B., Abtahi, H. R., Edalatifard, M., et al. (2024). Predation on bacterial pathogens by predatory bacteria of sewage origin: three days prey-predator interactions. BMC Microbiol. 24, 516. doi:10.1186/s12866-024-03672-z

Murphy, T. F., Brauer, A. L., Eschberger, K., Lobbins, P., Grove, L., Cai, X., et al. (2008). Pseudomonas aeruginosa in chronic obstructive pulmonary disease. Am. J. Respir. Crit. Care Med. 177 (8), 853–860. doi:10.1164/rccm.200709-1413OC

Naena, E. (2020). Yucca plant as treatment for Pseudomonas aeruginosa infection in Nile tilapia farms with emphasis on its effect on growth performance. Alex. J. Vet. Sci. 66, 64. doi:10.5455/ajvs.113537

National Center for Biotechnology Information (NCBI) (2022). Available at: https://www.ncbi.nlm.nih.gov/datasets/genome/GCF_027923905.1/

National Center for Biotechnology Information (NCBI) (2024). Available at: https://www.ncbi.nlm.nih.gov/datasets/genome/GCF_042660405.1/

Novick, R. P. (2021). Antibacterial particles and predatory bacteria as alternatives to antibacterial chemicals in the era of antibiotic resistance. Curr. Opin. Microbiol. 64, 109–116. doi: 10.1016/j.mib.2021.09.016

Orthova, I., Kämpfer, P., Glaeser, S. P., Kaden, R., and Busse, H. J. (2015). Massilia norwichensis sp. nov., isolated from an air sample. Int. J. Syst. Evol. Microbiol. 65, 56–64. doi:10.1099/ijs.0.068296-0

Ottaviani, D., Chierichetti, S., Angelico, G., Forte, C., Rocchegiani, E., Manuali, E., et al. (2018). Halobacteriovorax isolated from marine water of the Adriatic sea, Italy, as an effective predator of Vibrio parahaemolyticus, non-O1/O139 V. cholerae, V. vulnificus. J. Appl. Microbiol. 125 (4), 1199–1207. doi:10.1111/jam.14027

Ottaviani, D., Pieralisi, S., Rocchegiani, E., Latini, M., Leoni, F., Mosca, F., et al. (2020). Vibrio parahaemolyticus-specific Halobacteriovorax from seawater of a mussel harvesting area in the adriatic sea: Abundance, diversity, efficiency and relationship with the prey natural level. Front. Microbiol. 11, 1575. doi:10.3389/fmicb.2020.01575

Oyedara, O. O., Segura-Cabrera, A., Guo, X., Elufisan, T. O., Cantú González, R. A., and Rodríguez Pérez, M. A. (2018). Whole-genome sequencing and comparative genome analysis provided insight into the predatory features and genetic diversity of two Bdellovibrio species isolated from soil. Int. J. Genomics 2018, 1–10. doi:10.1155/2018/9402073

Pang, R., Xie, T., Wu, Q., Li, Y., Lei, T., Zhang, J., et al. (2019). Comparative genomic analysis reveals the potential risk of Vibrio parahaemolyticus isolated from ready-to-eat foods in China. Front. Microbiol. 10, 186. doi:10.3389/fmicb.2019.00186

Pepi, M., and Focardi, S. (2021). Antibiotic-resistant bacteria in aquaculture and climate change: A challenge for health in the mediterranean area. Int. J. Environ. Res. Public. Health 18 (11), 5723. doi:10.3390/ijerph18115723

Pérez, J., Moraleda‐Muñoz, A., Marcos‐Torres, F. J., and Muñoz‐Dorado, J. (2016). Bacterial predation: 75 years and counting! Environ. Microbiol. 18 (3), 766–779. doi:10.1111/1462-2920.13171

Ragunath, C., and Ramasubramanian, V. (2022). Dietary effect of Padina boergesenii on growth, immune response, and disease resistance against Pseudomonas aeruginosa in Cirrhinus mrigala. Appl. Biochem. Biotechnol. 194, 1881–1897. doi:10.1007/s12010-021-03770-y

Ribeiro, T. G., Izdebski, R., Urbanowicz, P., Carmeli, Y., Gniadkowski, M., and Peixe, L. (2021). Citrobacter telavivum sp. nov. with chromosomal mcr-9 from hospitalized patients. Eur. J. Clin. Microbiol. Infect. Dis. 40 (1), 123–131. doi:10.1007/s10096-020-04003-6

Rodgers, A. M., Lindsay, J., Monahan, A., Dubois, A. V., Faniyi, A. A., Plant, B. J., et al. (2023). Biologically relevant murine models of chronic Pseudomonas aeruginosa respiratory infection. Pathogens 12 (8), 1053. doi:10.3390/pathogens12081053

Roxo, I. C., Alarico, S., Fonseca, A., Machado, D., Maranha, A., Tiago, I., et al. (2025). Mycobacterium appelbergii sp. nov., a novel species isolated from a drinking water fountain in a rural community. Microorganisms 13, 1259. doi:10.3390/microorganisms13061259

Rubin, J., and Yu, V. L. (1988). Malignant external otitis: Insights into pathogenesis, clinical manifestations, diagnosis, and therapy. Am. J. Med. 85 (3), 391–398. doi:10.1016/0002-9343(88)90592-X

Seccareccia, I., Kost, C., and Nett, M. (2015). Quantitative analysis of Lysobacter predation. Appl. Environ. Microbiol. 81 (20), 7098–7105. doi:10.1128/AEM.01781-15

Sedláček, I., Holochová, P., Sedlář, K., Staňková, E., Šedo, O., Kralova, S., et al. (2025). Two new psychrotolerant Massilia species inhibit plant pathogens Clavibacter and Curtobacterium. Sci. Rep. 15, 26134. doi:10.1038/s41598-025-09734-0

Souza, C. F., Baldissera, M. D., Santos, R. C. V., Raffin, R. P., and Baldisserotto, B. (2017). Nanotechnology improves the therapeutic efficacy of Melaleuca alternifolia essential oil in experimentally infected Rhamdia quelen with Pseudomonas aeruginosa. Aquaculture 473 (20), 169–171. doi: 10.1016/j.aquaculture.2017.02.014

Speights, C. J., and McCoy, M. W. (2017). Range expansion of a fouling species indirectly impacts local species interactions. PeerJ 5, e3911. doi:10.7717/peerj.3911

Tao, L., Gao, D., Liu, Y., Sun, W., and Shan, X. (2026). The genus Aeromonas in aquaculture: A comprehensive review of prevalence, virulence, and antibiotic resistance with an emphasis on key pathogenic species. Rev. Aquac. 18 (1), e70097. doi:10.1111/raq.70097

Tatusov, R. L., Galperin, M. Y., Natale, D. A., and Koonin, E. V. (2000). The COG database: a tool for genome-scale analysis of protein functions and evolution. Nucleic Acids Res., 28(1), 33–36. doi:10.1093/nar/28.1.33

Thanigaivel, S., Thomas, J., Vickram, A. S., Gulothungan, G., Nanmaran, R., and Jenila Rani, D. (2023). Antioxidant and antibacterial efficacy of Chaetomorpha linum and its toxicological evaluation for the prophylactic treatment against Pseudomonas aeruginosa infection in Labeo rohita. J. Appl. Aquac. 35, 350–369. doi: 10.1080/10454438.2021.1961966

Tian, M., Jia, P., Wu, Y., Yu, X., Wu, S., Yang, L., et al. (2023). Diversity and distribution characteristics of culturable bacteria in burqin glacier No. 18, altay mountains, China. Diversity 15 (9), 997. doi: 10.3390/d15090997

Triyaningsih, Sarjito, and Prayitno, S. B. (2014). Patogenisitas Aeromonas hydrophila yang diisolasi dari lele dumbo (Clarias gariepinus) yang berasal dari Boyolali. J. Aquac. Manag. Technol. 3 (2), 11–17.

Turnip, E. R., Widanarni, W., and Meryandini, A. (2018). Selection of lactic acid bacteria as a probiotic and evaluated its performance on gnotobiotic catfish Clarias sp. J. Akuakultur Indones. 17 (1), 68–80. doi:10.19027/jai.17.1.68-80

Verschuere, L., Rombaut, G., Sorgeloos, P., and Verstraete, W. (2000). Probiotic bacteria as biological control agents in aquaculture. Microbiol. Mol. Biol. Rev. 64 (4), 655–671. doi: 10.1128/MMBR.64.4.655-671.2000

Wang, H., Lou, J., Gu, H., Luo, X., Yang, L., Wu, L., et al. (2016). Efficient biodegradation of phenanthrene by a novel strain Massilia sp. WF1 isolated from a PAH-contaminated soil. Environ. Sci. Pollut. Res. 23, 13378–13388. doi: 10.1007/s11356-016-6515-6

Wang, S. (2020). Isolation, polyphasic taxonomy and predatory characteristics of Bradymonabacteria. Jinan: Shandong University. 10.27272/d.cnki.gshdu.2020.000367.

Wang, Y., Liang, Q., Lu, B., Shen, H., Liu, S., Shi, Y., et al. (2021). Whole-genome analysis of probiotic product isolates reveals the presence of genes related to antimicrobial resistance, virulence factors, and toxic metabolites, posing potential health risks. BMC Genomics 22 (1), 210. doi: 10.1186/s12864-021-07539-9

Weon, H. Y., Yoo, S. H., Kim, S. J., Kim, Y. S., Anandham, R., and Kwon, S. W. (2010). Massilia jejuensis sp. nov. and Naxibacter suwonensis sp. nov., isolated from air samples. Int. J. Syst. Evol. Microbiol. 60(8), 1938–1943. doi: 10.1099/ijs.0.015479-0

Wingender, J., Strathmann, M., Rode, A., Leis, A., and Flemming, H. C. (2001). Isolation and biochemical characterization of extracellular polymeric substances from Pseudomonas aeruginosa. Method. Enzymol. 336, 302–314. doi: 10.1016/S0076-6879(01)36597-7

Xi, Y., Pan, Y., Li, M., Zeng, Q., and Wang, M. (2024). Evaluation of the application potential of Bdellovibrio sp. YBD-1 isolated from Yak faeces. Sci. Rep. 14, 13010. doi: 10.1038/s41598-024-63418-9

Xie, Z., and Tang, H. (2017). ISEScan: automated identification of insertion sequence elements in prokaryotic genomes. Bioinformatics 33(21), 3340–3347. doi:10.1093/bioinformatics/btx433

Xu, A., Liu, C., Zhao, S., Song, Z., and Sun, H. (2023). Dynamic distribution of Massilia spp. in sewage, substrate, plant rhizosphere/phyllosphere and air of constructed wetland ecosystem. Front. Microbiol. 14, 1211649. doi: 10.3389/fmicb.2023.1211649

Xu, J., Feng, G., and Yan, Y. (2025). Effects of polystyrene nanoplastics and copper on gill tissue structure, metabolism, and immune function of the Chinese mitten crab (Eriocheir sinensis). Front. Mar. Sci. 12, 1538734. doi:10.3389/fmars.2025.1538734

Xu, X. H., Zhang, Y. Q., Yan, B. L., Xu, J. T., Tang, Y., and Du, D. D. (2014). Immunological and histological responses to sulfide in the crab Charybdis japonica. Aquat. Toxicol. 150, 144–150. doi:10.1016/j.aquatox.2014.03.006

Yang, Z., Zhang, Y., Hu, K., Liu, L., Cai, H., Zhang, F., et al. (2018). Etiological and histopathological study on hepatopancreatic necrosis syndrome in Eriocheir sinensis. Acta Hydrobiol. Sin. 42 (1), 17–25. doi:10.7541/2018.003

Zago, V. (2020). Marine microorganisms producing carbapenemases and β-lactamases as environmental reservoirs of antibiotic resistance genes constituting a risk for human health. Italy: University of Verona.

Zhang, C., Shi, R., Yuan, Z., Zhang, D., Wang, Z., Luo, F., et al. (2025). Isolation, identification, pathogenicity and development of a rapid detection technique of Pseudomonas aeruginosa strain GX2021 causing bacterial shell erosion disease in farmed Bellamya aeruginosa. J. Invertebr. Pathol. 212, 108375. doi:10.1016/j.jip.2025.108375

Zhang, J., Wang, M., Hu, L., Zhang, Q., Chen, E., Wang, Z., et al. (2022). A universal coating strategy for inhibiting the growth of bacteria on materials surfaces. Front. Chem. 10, 1043353. doi:10.3389/fchem.2022.1043353

Zhang, Y. Y., Huang, Y. F., Liang, J., and Zhou, H. (2022). Improved up-and-down procedure for acute toxicity measurement with reliable LD_50_ verified by typical toxic alkaloids and modified Karber method. BMC Pharmacol. Toxico. 23(1), 3. doi:10.1186/s40360-021-00541-7

Zhou, J. S., Shu, Q., Rutherfurd, K. J., Prasad, J., Gopal, P. K., and Gill, H. S. (2000). Acute oral toxicity and bacterial translocation studies on potentially probiotic strains of lactic acid bacteria. Food Chem. Toxicol. 38 (2-3), 153–161. doi:10.1016/S0278-6915(99)00154-4

